# *ΔF508-Cftr* mutation in genetically diverse Collaborative Cross mice yields novel disease-relevant phenotypes for cystic fibrosis

**DOI:** 10.1101/2023.01.31.526098

**Authors:** Barbara Sipione, Nicola Ivan Lorè, Francesca Sanvito, Giacomo Rossi, Alessia Neri, Fabrizio Gianferro, Anna Sofia Tascini, Alessandra Livraghi-Butrico, Cristina Cigana, Alessandra Bragonzi

## Abstract

Mutations of *Cystic Fibrosis Transmembrane conductance Regulator* (*CFTR*) lead to Cystic Fibrosis (CF), but the substantial phenotypic variations are determined by non-*CFTR* allelic diversity. To map novel disease phenotypes in a CF mouse model, we used Collaborative Cross (CC) mice, a highly genetically diverse mouse resource population. *ΔF508-Cftr* homozygosity produced a fully penetrant lethal phenotype by eight weeks in two CC lines. The lethality of CC006*ΔF508/ΔF508* was fully prenatal while CC037*ΔF508/ΔF508* showed either prenatal or postnatal lethality. Novel phenotypes of CC037*ΔF508/ΔF508* were revealed early in life including respiratory and systemic inflammatory profiles, and blood, bone marrow, pancreas, heart, and reproductive tract pathologies. Severe intestinal blockage was observed as common in other CF mouse models. These results suggest that the exploration of CF disease phenotypes in a mouse population with diverse genetic profiles is needed to map the genetic origin of currently unidentified disease traits and their potential translation to humans.

## Introduction

Cystic fibrosis (CF) is an autosomal recessive disease caused by mutations in the *cystic fibrosis transmembrane conductance regulator* (*CFTR*) gene ^1–3^. The *CFTR* gene encodes a cAMP-responsive chloride channel expressed at the surface of most epithelial cells that can also affect the transport of other ions and water. CF affects several organs including the pancreas, intestine, liver, gall bladder, sweat glands, and male reproductive tract, but the lung is where fatal complications occur. Absence of CFTR-mediated chloride secretion has been functionally linked to airway surface dehydration, which leads to the accumulation of thick mucus, increased susceptibility to infection by several pathogens, and inflammation ^4–6^. Although CF is classified as a single-gene disorder, patients who carry the same *CFTR* mutations exhibit substantial phenotypic variation, with particular relevance to lung disease, of which >50% is explained by non-*CFTR* genetic variation ^3,7^. Although the identification of non-*CFTR* genetic variations has guided research in human studies, there continues to be a substantial gap in our understanding of how host genetics influence CF disease severity.

The 13 CF murine models and models in seven additional species (e.g. pig, ferret, rat, zebrafish, sheep, drosophila and rabbit), established so far, have contributed at different levels to our understanding of CF multiorgan disease, but only a few would be amenable to conduct large scale genotype/phenotype association studies ^8,9^. Murine models present major challenges related to the limited number of genetically-redundant inbred strains, mainly C57BL/6, used for their generation. This constraint limits allelic diversity and likely explains the failure of these models to recapitulate the broad spectrum of clinical conditions observed in people with CF (pwCF). Progress towards the development of other *CFTR*-targeted small animal models has been made in ferrets, rats, and rabbits ^10–12^, and these laboratory models recapitulate some aspects of human disease but fail to recapitulate the polymorphic variations that are hallmarks of the human population. Large animal models of CF, including pig and sheep ^13,14^, have demonstrated good suitability for translation of research findings to humans, but cost and practical considerations limit their wide-spread availability for investigating disease mechanisms and treatments. The inability to investigate genetic variability in an informative yet economically feasible way remains a major concern.

The Collaborative Cross (CC) mouse population ^15^ represents an opportunity to mimic the genetic diversity of the human population. CC mice carry high allelic diversity due to systematic outcrosses of eight founder strains. Five of the eight founders are common laboratory strains (A/J, C57BL/6J, 129S1/SvImJ, NOD/LtJ, and NZO/HiLtJ), and three are wild-derived strains (CAST/Ei, PWK/PhJ, and WSB/EiJ) ^16^. Consequently, CC mice have greater recombination and genetic variation (90%) than other reference panels (30%), representing a genetic diversity twice as large as that of the human population ^17^. A spectrum of pathogenic phenotypes, never previously reported for classical inbred strains, has been observed in CC mice, which have been studied for a wide range of biomedically relevant traits. CC mice have been used extensively to study host-pathogen interactions ^18^, including viral, bacterial and fungal infection, and several diseases affected by complex traits (e.g. metabolic and neurological disorders, cancer, behavior) ^19^, but they have never been used to test the influence of genetic background on phenotypes elicited by a specific genetic mutation.

Here, we addressed how we can incorporate genetic variability to generate a diverse mouse model for CF and fill the gap between the CF manifestations in patients and mouse models of this disease. To this end, we introduced the Δ*F508-Cftr* mutation in two CC lines (CC006 and CC037) selected for their high allelic diversity and phenotypic traits relevant for CF disease, including susceptibility to *Pseudomonas aeruginosa* infection ^20,21^. In addition to previously described phenotypes, CC mice carrying the *ΔF508-Cftr* mutation exhibited novel pathological phenotypes with multiple organ involvement and open new hypotheses regarding the manifestation of CF disease in humans.

## Results

### The ΔF508-Cftr mutation in CC lines caused prenatal and postnatal lethality

To cover high genetic and phenotypic variability relevant to CF disease, we ranked 39 genetically diverse CC lines for the different responses to *P. aeruginosa* infection and other respiratory pathogens (**Table S1**). Among 11 CC lines susceptible to *P. aeruginosa* infection ^20,21^, two lines were also susceptible to *Klebsiella pneumonia* and *Aspergillus fumigatus* ^22,23^; while 11 CC lines were resistant to infection. Next, we excluded murine strains with spontaneous pathological phenotypes and those not characterized for genotypic and phenotypic traits (http://csbio.unc.edu/CCstatus). Strains with the highest number of founder strains’ genotypes and the lowest heterozygosity were considered favorable for selection. Good fertility was also considered an essential trait, considering the reproductive difficulties of CC lines and CF mice. Based on these criteria and the score assigned, CC006 and CC037 lines were selected for insertion of the *ΔF508-Cftr* mutation by high-speed backcrossing with C57BL/6J (Cftr^tm1Kth^) mice ^24^ (**Fig. S1**). The heterozygous progeny of *ΔF508-Cftr* mice in both lines appeared normal and were fertile, although the CC037 line produced more pups per litter (mean litter size of 5.3 pups) than the CC006 line (mean litter size of 3.5 pups) (**Table S2**).

Breeding the heterozygous *CC037ΔF508/wt* mice produced 816 *wt/wt* mice (29%), 1655 *ΔF508/wt* (58%), and 369 *ΔF508/ΔF508* (13%), a percentage significantly different from the predicted Mendelian ratio (p< 0.00001) (**Fig. 1A** and **Table S2**) and suggesting 12% of prenatal lethality for homozygous *ΔF508-Cftr* in CC037 mice. Postnatal mortality was observed during the first five days after birth (29%), declined in the second and third week (17%), and peacked again between the third and fourth week of life before weaning (33%) (**Fig. 1A** and **Table S3**). 21% of the mice died post-weaning before 8 weeks. In contrast, CC037*ΔF508/wt* and *wt/wt* show low postnatal mortality at 6% and 4% respectively, by the first five days of life. The overall survival curve showed major significant differences between CC037*ΔF508/ΔF508* and *wt/wt* (p<0.0001), regardless of sex (**Fig. 1B**).

**Fig. 1.**
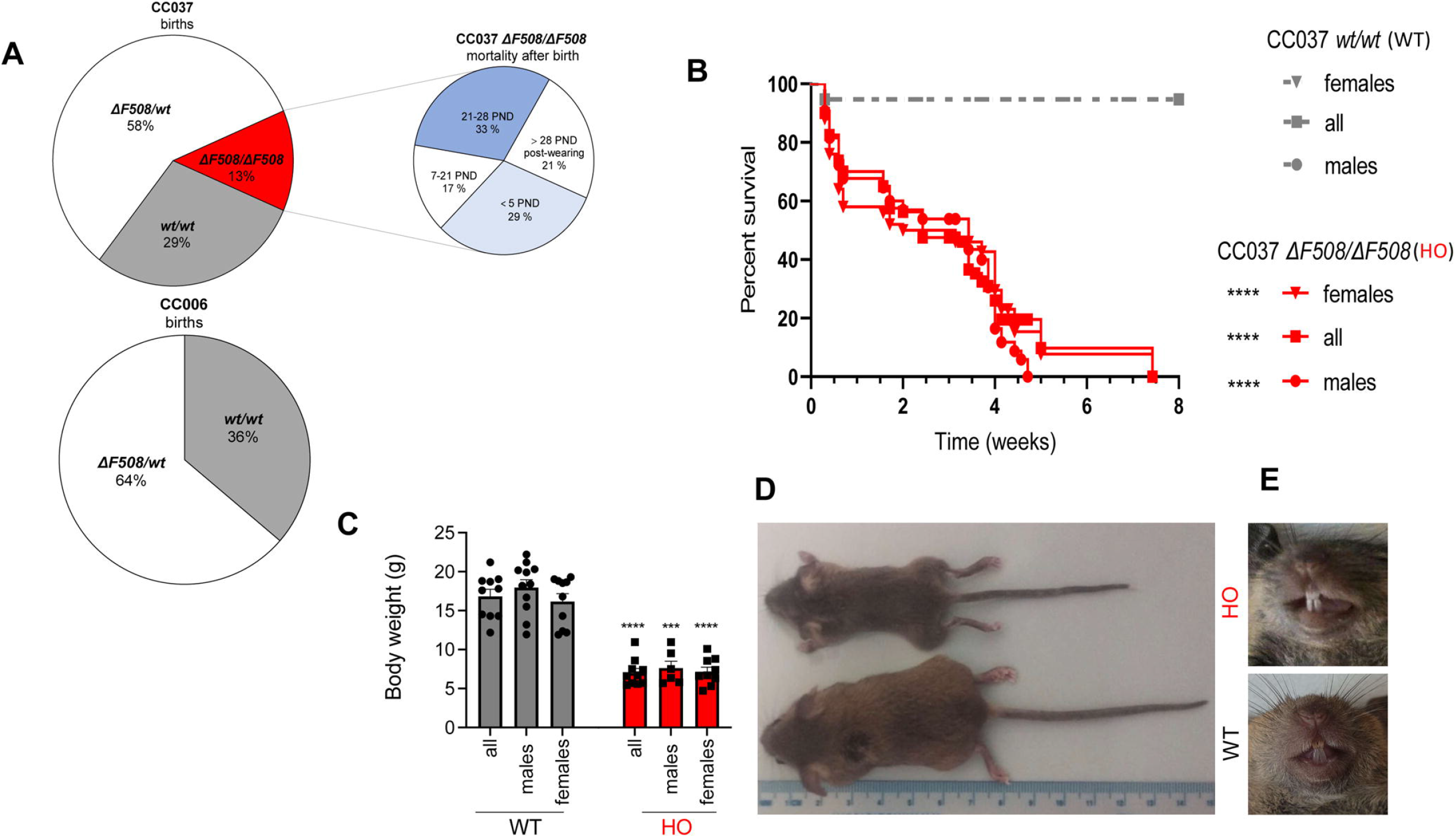
Survival and growth of CC037 and CC006 *ΔF508/ΔF508* compared to *wt/wt* and *ΔF508/wt* mice. **A**) Percentage of births of three different genotypes with respect to murine *Cftr (wt/wt, ΔF508/wt* and *ΔF508/ΔF508)* obtained by intercrossing heterozygous *CC037ΔF508/wt* and CC006*ΔF508/wt* mice. Next, mortality of CC037*ΔF508/ΔF508* is indicated at different post-natal day (PND). Weaning occurred at PND28. **B**) Survival curves for all mice, including female and male, *CC037wt/wt* (WT) (n*=*153) and CC037*ΔF508/ΔF508* (HO) mice (n*=*80). The data are pooled from the offspring of different breeding pairs (n males: *wt/wt=* 70, *ΔF508/ΔF508=37;* n females: *wt/wt=83, ΔF508/ΔF508=43)* Statistical significance between *wt/wt* and *ΔF508/ΔF508*, male and female, determined by Mantel Cox test is indicated as follows: ****, *P*<0.0001. **C**) Differences in body weight between all CC037*wt/wt* (n=21) and *ΔF508/ΔF508* (n=15) mice and between male and female *CC037wt/wt* and *ΔF508/ΔF508* at PND 28 (n males: *wt/wt=11, ΔF508/ΔF508=6; n* females: *wt/wt*=10*, ΔF508/ΔF508=9*). Statistical significance between *ΔF508/ΔF508* and *wt/wt* determined by Mann Whitney test is indicated as follows: ***, *P*<0.001; ****, *P*<0.0001. **D**) Representative mice at PND28, with the genotype indicated. **E**) Dentition in CC037*ΔF508/ΔF508* compared to *wt/wt* mice.

Compared to *CC037wt/wt* littermates, *ΔF508/ΔF508* mice showed significantly reduced body weight (p<0.0001) (**Fig.1C, D**) at postnatal day (PND) 28. No significant differences between males and females were observed for both mortality and body weight. Similar to other CF mouse models, CC037*ΔF508/ΔF508* exhibited white, chalky enamel that differed from the yellow-brown, smooth tooth enamel in *wt/wt* mice (**Fig. 1E**).

Breeding heterozygous CC006*ΔF508/wt* mice produced 99 *wt/wt* mice (36%),174 *ΔF508/wt* (64%), and no *ΔF508/ΔF508* (0%), a percentage significantly different from the predicted Mendelian ratio (p< 0.00001), suggesting that the homozygous mutation in the CC006 genetic context resulted in fully penetrant prenatal lethality (**Fig. 1A, Table S2**). CC006 *ΔF508/wt* and *wt/wt* mice showed low postnatal mortality at 9% and 4% respectively, by the first five days of life (**Table S3**).

These results show that the same *ΔF508-Cftr* mutation in the context of different genetic backgrounds influences prenatal and postnatal survival.

### Sequencing of CC037 and CC006 lines with the ΔF508 mutation showed different founder contributions to the genomes

CC037 and CC006 mice carrying *ΔF508-Cftr* mutation after high-speed backcrossing were genotyped and compared with the original line. Whole genome sequencing (WGS) showed that the genome of *CC037ΔF508/wt* line after backcross has haplotype contributions from six of the eight founders with no major differences from the CC037 line originally described ^17^. Three founders (NOD/ShiLtJ, C57BL/6J, and A/J) were more represented, three were less represented (129S1/SvlmJ, CAST/EiJ, and WSB/EiJ) and two were absent (NZO/H1LtJ and PWK/phJ) (**Fig. 2A and B** and **Table 1**). We found 5,006,279 variants (SNPs) in the CC037 *ΔF508/wt* with 1,159,506 InDels, using C57BL6/J as a reference genome. After high-speed backcrossing, the percentage of autosomal heterozygosity in the CC037 *ΔF508/wt* strain was increased to 3.72% compared to 0.02% in the original CC037 line^17^.

**Fig. 2.**
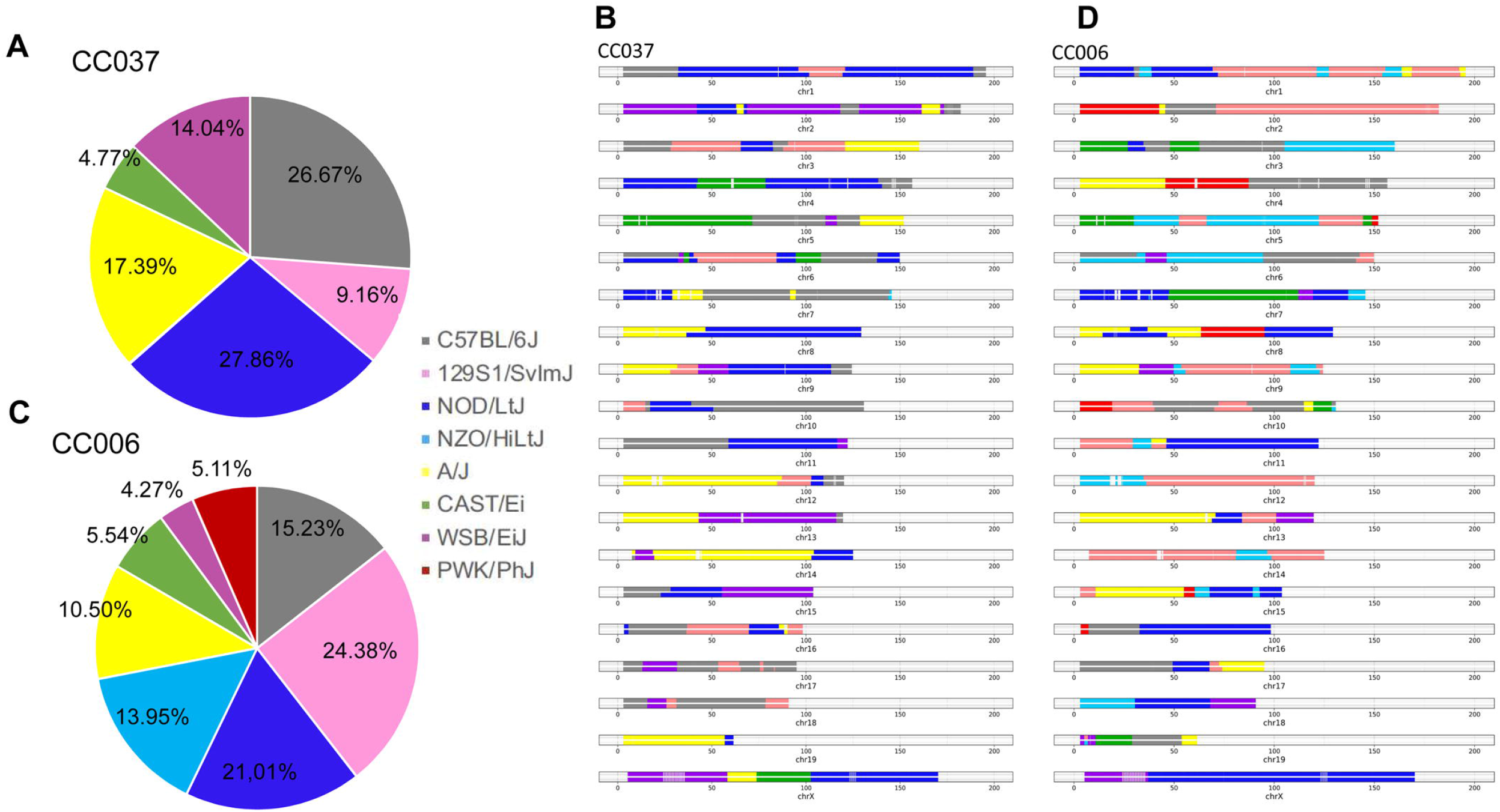
The CC037 and CC006 genome and their *Cftr*. Haplotype structure for the sequenced representative of the *CC037ΔF508/wt* (**A, B**) and *CC006ΔF508/wt* (**C, D**) strains with founder contribution to each chromosome and percentage to the genomes. The following colors were used to represent the eight founders strains of the CC: NOD/ShiLtJ (dark blue), C57BL/6J (gray), A/J (yellow), 129S1/SvlmJ (pink), CAST/EiJ (green), WSB/EiJ (purple), NZO/H1LtJ (light blue) and PWK/PhJ (red).

**Table 1.** Genomic information for CC006 and CC037 lines with *ΔF508* mutation sequenced murine strains.

| Strain |  |  | % Heterozygosity | Haplotype contributions |  |  | *Cftr* |  | InDels |
| --- | --- | --- | --- | --- | --- | --- | --- | --- | --- |
|  | Total SNPs | Total InDels |  | More-represented | Less-represented | Absent | strain contribution | SNPs |  |
| CC037<br>$\Delta F508/wt$ | 5,006,279 | 1,159,506 | 3.72% | NOD/ShiLtJ (27.86%)<br>C57BL/6J (26,67%)<br>A/J (17,39%) | *WSB/EiJ (14.04%)<br>129S1/SvImJ (9.16%)<br>**CAST/EiJ (4.77%) | NZO/H1LtJ<br>PWK/phJ | NOD/ShiLtJ<br>+<br>C57BL/6J<br>(*Cftr*<sup>tm1Kth</sup>) | 1,253 | 248 |
| CC006<br>$\Delta F508/wt$ | 6,839,626 | 1,486,595 | 6.48% | 129S1/SvImJ (24,38%)<br>NOD/ShiLtJ (21,01%)<br>C57BL/6J (15,23%)<br>NZO/H1Lt (13,95%) | A/J (10,50%)<br>**CAST/EiJ (5,54%)<br>**PWK/PhJ (5,11%)<br>*WSB/EiJ (4,27%) | - | NZO/H1LtJ<br>+<br>C57BL/6J<br>(*Cftr*<sup>tm1Kth</sup>) | 1,350 | 291 |

\* indicate wild derived strains that represent three *Mus musculus* subspecies.
\*\* indicates strains of non-*domesticus* origin.

The CC006 *ΔF508/wt* line had haplotype contributions from eight founders with no major differences from the line CC006 originally described^17^. Four founders were more represented (C57BL/6J, 129S1/SvlmJ, NOD/ShiLtJ, and NZO/H1LtJ) and four were less represented (A/J, CAST/EiJ, WSB/EiJ, and PWK/phJ) (**Fig. 2C and D** and **Table 1**). We found 6,839,626 SNPs in the CC006 *ΔF508/wt* with 1,486,595 InDels, using C57BL6/J as a reference genome. The percentage of heterozygosity in the autosomes of the CC006 *ΔF508/wt* strain was 6.48% compared to 4.12% in the original CC006 line^17^.

Thus, our CF mouse models have substantial genomic differences from those previously reported with the *Cftr* mutation in the C57BL/6J background and share less than a quarter of the genome with the latter. Interestingly, haplotype reconstruction analysis shows that the chr6 in both lines has a haplotype block including the *Cftr* locus in heterozygosity (**Fig2 B and D**). Specifically, for the CC037 strain, the haplotype block has a length of 33Mb and is characterized by heterozygosis of the NOD/ShiLtJ founder haplotype and the C57BL/6J (Cftr^tm1Kth^) founder haplotype. The CC006 haplotype block measures 31Mb and contains the NZO/H1LtJ and C57BL/6J (Cftr^tm1Kth^) haplotypes.

### CC037ΔF508/ΔF508 mice showed severe gut obstruction and increased gastric mucins

Gross observation and histological evaluation indicated that CC037*ΔF508/ΔF508* mice suffer from gut obstruction with failure to pass stool (**Fig. 3A**) while no gross pathology of the gut was observed in *wt/wt* mice. The gut obstruction in CC037*ΔF508/ΔF508* mice was associated with intestinal luminal mucus accumulation and mucous cell hyperplasia (**Fig. 3B-D**). Interestingly, enhanced production of mucus that was sticky and adhered to the intestinal wall was observed in CC037*ΔF508/ΔF508* mice, while in *wt/wt* congenic mice the mucus was present in a lower amount, and it was non-adherent. (**Fig. 3E-G**). These data were similar to those obtained in other CF mouse models and support the strong influence of *ΔF508-Cftr* mutation on intestinal phenotypes, regardless of genetic background.

**Fig. 3.**
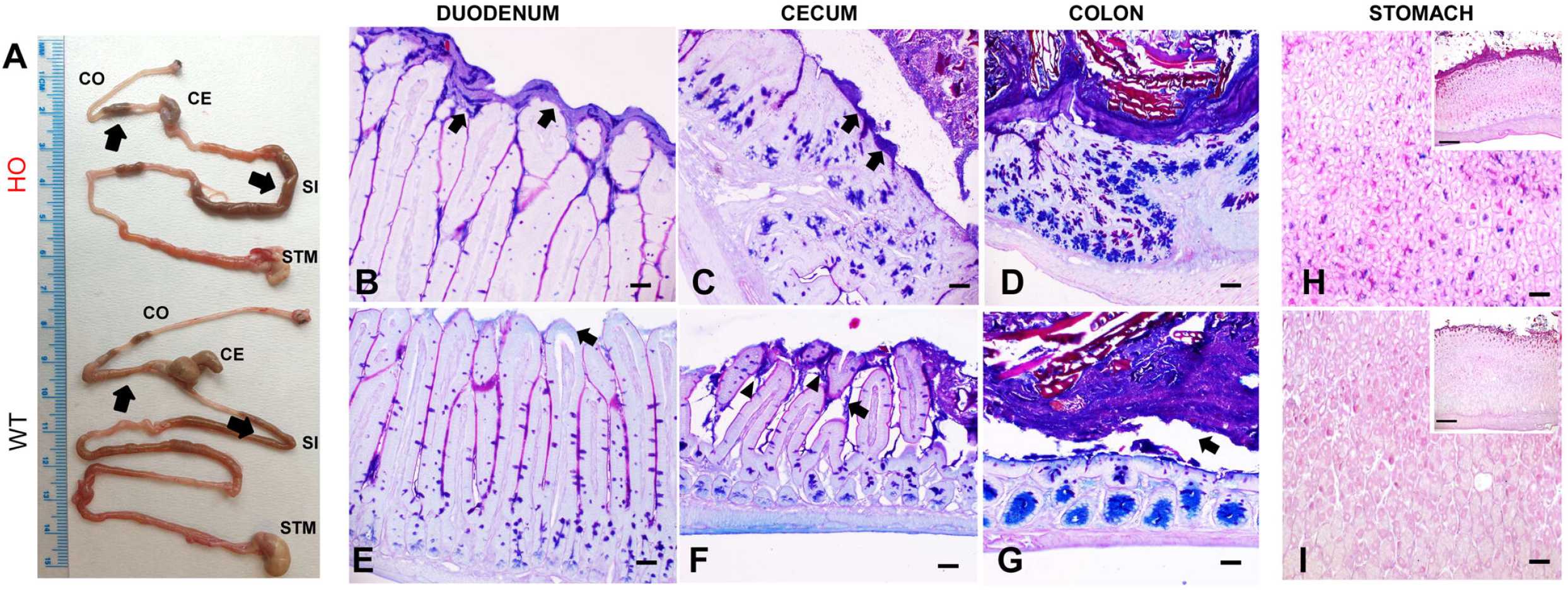
Gut and stomach disease phenotype in CC037*ΔF508/ΔF508* compared to *wt/wt* mice. **A**) Representative pictures of intestines at PND28. Arrows indicate the location of obstructions in the small and large intestines of CC037*ΔF508/ΔF508* (HO) mice and the absence of obstruction in *wt/wt* (WT) ones. **B-G**) Gut histology with AB-PAS staining at PND28. Arrows indicate enhanced production of mucus adhering to epithelial cells in HO mice (**B-D**) and normal secretion of mucus that did not adhere to epithelial cells in WT mice (**E-G**). **H, I**) Stomach histochemistry of HO mice showing increased secretion of mucins, enriched of acidic and alcianophyl blue stained mucins, revealed by AB-PAS staining compared with WT mice. Scale bars= 100 μm (B-D and E-G), 150 μm (H-I), and 250 μm (inserts in H-I). *n*=14 WT and *n*=15 HO mice (number of males: WT=7, HO=8; number of females: WT=7, HO=7). STM: stomach; SI: small intestine; CE: cecum; CO: colon.

In the stomach, elevated production of gastric mucins in the mucosa was observed in CC037 *ΔF508/ΔF508* mice, with differences in mucus quantity and typology as compared to *wt/wt* mice (**Fig. 3H-I**). Specifically, in the deeper area of the mucosa, the mucous glands in CC037*ΔF508/ΔF508* mice showed increased prevalence of acidic mucins (alcian blue^+^-AB^+^) compared with more neutral mucins (periodic acid Shift^+^ - PAS^+^) of *wt/wt* mouse (**Fig. 3H-I**, inserts).

### CC037ΔF508/ΔF508 mice exhibited an early respiratory inflammatory profile in the lung

The lungs of CC037*ΔF508/ΔF508* mice were significantly smaller in size than those of *wt/wt* mice (**Fig. 4A** and **Fig. S2A**). However, the ratio of the lung weight to body weight of CC037*ΔF508/ΔF508* mice was significantly higher compared to *wt/wt* (**Fig. 4B**) and the quantification of lung protein content showed significantly higher levels for CC037*ΔF508/ΔF508* compared to *wt/wt* mice indicating possible pathological changes (**Fig. 4C**).

**Fig. 4.**
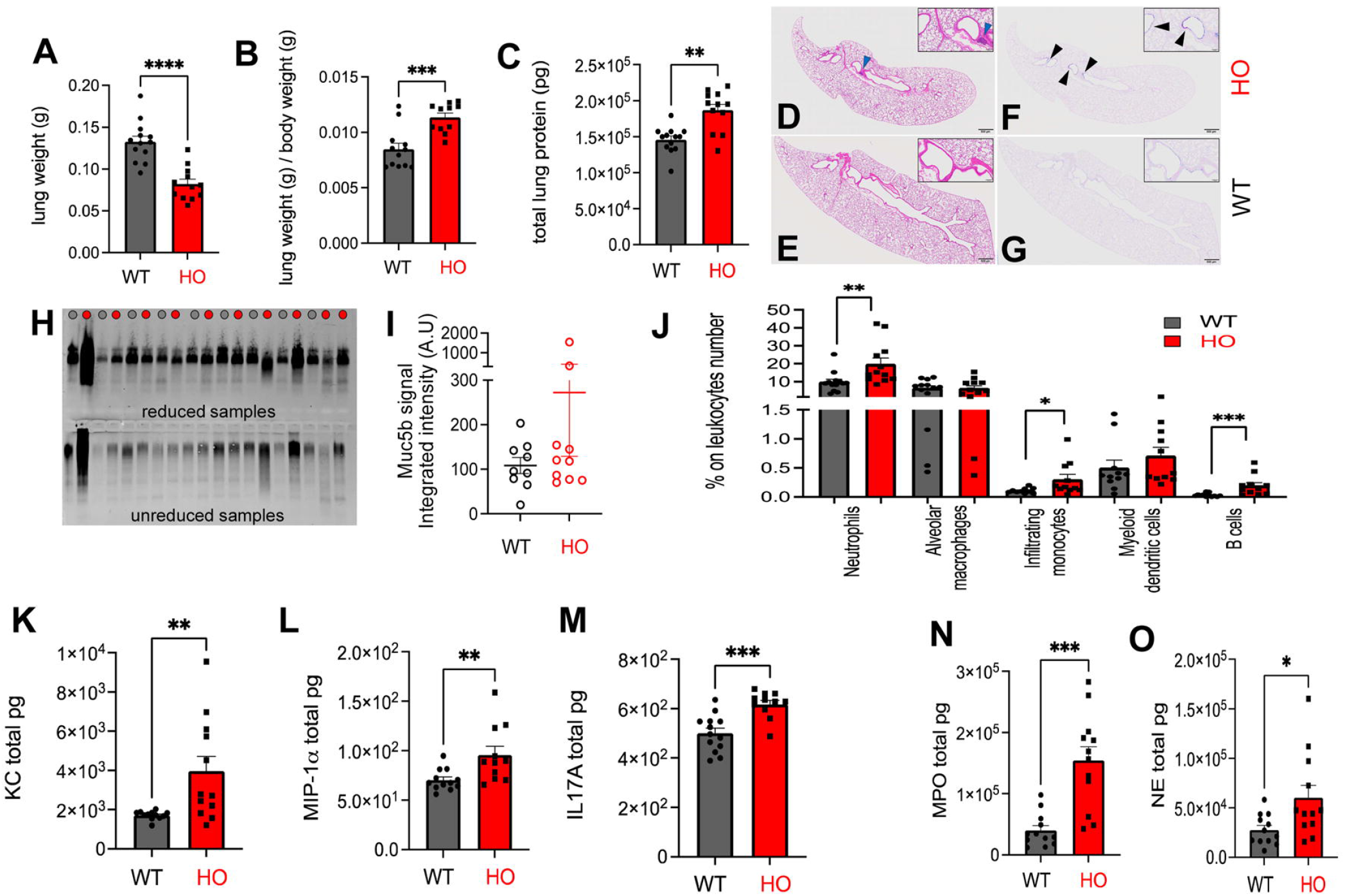
Respiratory disease phenotype of CC037*ΔF508/ΔF508* compared to *wt/wt* mice. **A)** Weight (g) of lungs and **B**) lung weight (g) normalized to body weight (g) from CC037*ΔF508/ΔF508* (HO) compared to *wt/wt* (WT) mice at PND28. **C**) Total proteins (pg) in lung lysates were evaluated by Bradford assay in HO compared to WT mice. **D-G**) Representative lung histology H&E (D-E) and AB-PAS (F-G) staining in HO and WT mice showing absence of gross morphological changes. Scale bars= 500 μm, insets 100 μm. Focal mucous cell metaplasia (black harrowheads) and submucosal lymphocytic aggregates (blue arrowhead) were occasionally observed in HO mice. *n*=14 WT and *n*=15 HO mice (number of males: *wt/wt=7*, HO=8; number of females: WT =7, HO=7). **H-I**) Mucin agarose western blot of BAL harvested from PND28 HO mice (red dot) and WT littermates (gray dot). Samples were run in reduced (upper panel) and unreduced (lower panel) form (**H**) to assess overall abundance and polymeric assembly, respectively. Densitometric analysis of Muc5b specific signal (**I**) was performed on reduced samples. **J**) Percentage of neutrophils, alveolar macrophages, infiltrating monocytes, myeloid dendritic cells and B cells on CD45+ cells measured by FACS analysis in lungs of HO and WT mice collected at PND28. **K-M**) Pg of KC, MIP-1α and IL-17A normalized for lung weight (g) measured by Bio-Plex assay in lung supernatants after physical dissociation. **N**) Pg of MPO and **O**) NE normalized for lung weight (g) measured in lung supernatants after physical dissociation. Bars represent mean ± standard error of the mean pooled from *n*=13 WT and *n*=12 HO mice pooled from 9 independent experiments. Statistical significance determined by Mann-Whitney test is indicated as follows: *, p<0.05; **, p<0.01; ***, p<0.001.

At PND 28, gross observation and histopathological analysis did not reveal major phenotypic changes in most of the CC037*ΔF508/ΔF508* lung compared to *wt/wt* (**Fig. 4D-E**). However, focal inflammatory infiltration, mucous cell metaplasia, and luminal mucus accumulation were sporadically observed in the bronchi (**Fig. 4F**), trachea and nasal cavity *CC037ΔF508/ΔF508* (**Fig. S2 B-E**) compared to *wt/wt* (**Fig. 4G, Fig. S2 F-I**). Mucin agarose western blot analysis of BAL isolated from CC037*ΔF508/ΔF508* and *wt/wt* littermates at PND28 confirmed the occasional presence of high Muc5b levels in CC037*ΔF508/ΔF508* mice (**Fig. 4H, upper panel, Fig. 4I**). Muc5b polymerization was similar in CC037*ΔF508/ΔF508* mice and WT littermates, as indicated by the presence of high molecular weight molecules in unreduced samples (**Fig. 4H, lower panel**). To define whether *ΔF508-Cftr* mutation can determine early inflammation, we performed FACS-assisted immunophenotyping of lung cells infiltrates in the lung. At PND 28, percentage of neutrophils and, to a less extent, infiltrating monocytes and B cells were significantly higher in the parenchyma of *CC037ΔF508/ΔF508* than in *wt/wt* mice (**Fig. 4J**). Cytokine/chemokine profiles indicated that keratinocyte chemoattractant (KC), macrophage inflammatory protein-1α (MIP-1α) and interleukin 17A (IL-17A) were significantly increased in CC037*ΔF508/ΔF508* compared to *wt/wt* mice (**Fig. 4K-M**). Other cytokines/chemokines and growth factors were not significantly different or not detected (**Table S4**). Furthermore, myeloperoxidase (MPO) and neutrophil elastase (NE) were significantly increased in CC037*ΔF508/ΔF508* compared to *wt/wt* mice suggesting ongoing inflammation (**Fig. 4N** and **O**). Quantitative data support the notion that *ΔF508-Cftr* mutation in the context of CC037 genetic background can elicit an early-onset inflammatory profile in the lung.

### Systemic inflammation and microcytic erythrocytosis were evidenced in CC037ΔF508/ΔF508 mice

Next, hematological analyses were performed to assess systemic inflammation and associated pathologies at PND28. Total counts of white blood cells were similar in *CC037ΔF508/ΔF508* and *wt/wt* mice, but the percentage of specific populations differed (**Table S5**). CC037*ΔF508/ΔF508* mice showed a significantly higher percentage of neutrophils than *wt/wt* mice (**Fig. 5A**). Moreover, a significantly lower lymphocyte percentage was observed in CC037*ΔF508/ΔF508* mice, but no differences were observed in monocyte, eosinophils, and basophils levels. These data suggest that *ΔF508-Cftr* mutation determines systemic inflammation early in life. Furthermore, red blood cell count (RBC), mean corpuscular volume (MCV) and mean corpuscular hemoglobin (MCH) differed significantly in the two murine genotypes (**Fig. 5B-D**). Levels of RBC were significantly increased in CC037*ΔF508/ΔF508* compared to the *wt/wt*, indicating an increased total amount of red blood cells. A significant reduction in MCV and MCH of CC037*ΔF508/ΔF508* compared to *wt/wt* indicates reduced red blood cells size and hemoglobin content in HO mice. Other RBC and hemoglobin parameters were not significantly different (**Table S5**). Overall, hematological analysis suggested microcytic erythrocytosis.

**Fig. 5:**
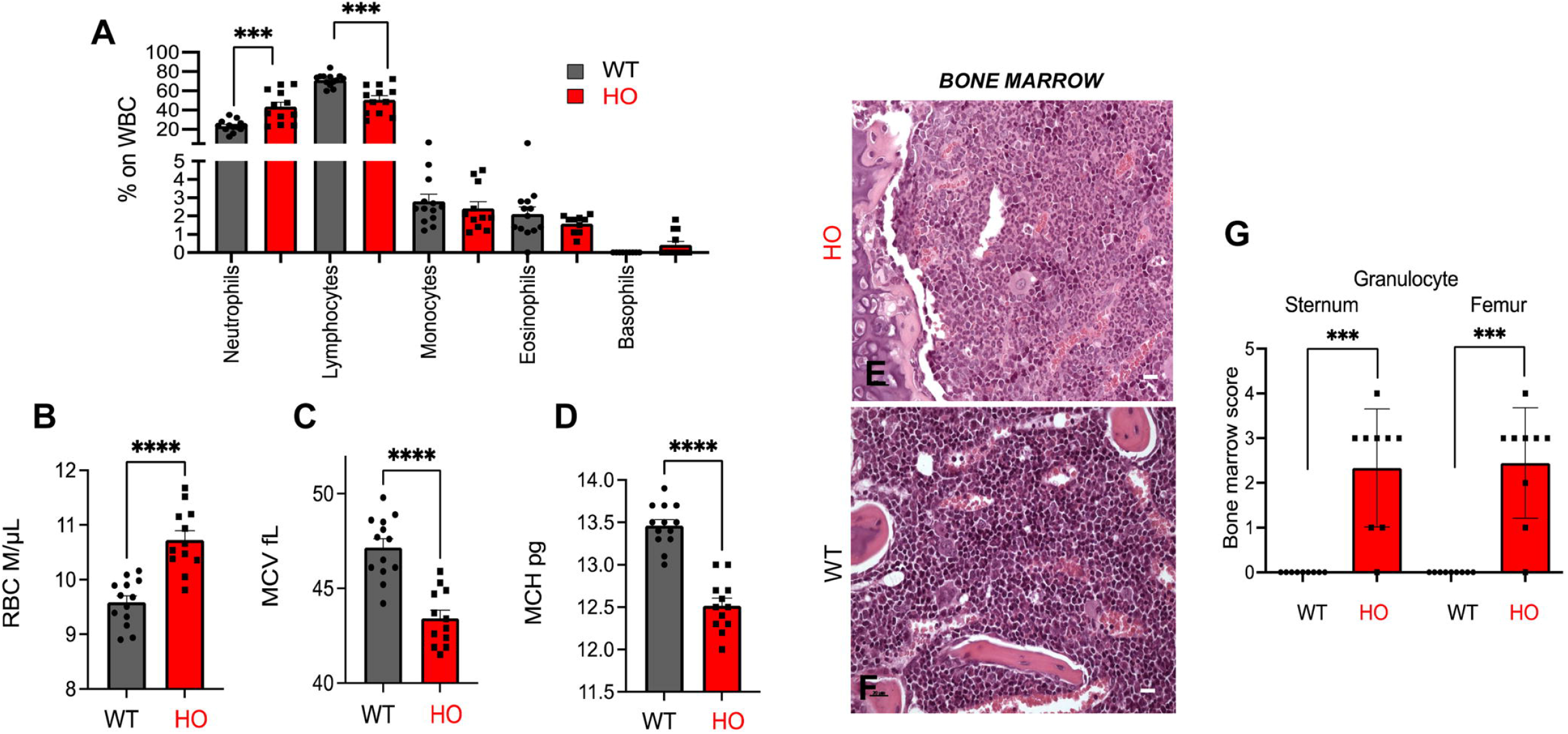
Disease phenotypes in the blood and bone marrow of *CC037ΔF508/Δ F508* and *CC037wt/wt* mice. **A**) Haematological analysis with percentage of specific populations on white blood cells (WBC) and (**B**) Red Blood Cell count (RBC), (**C**) Mean Corpuscular Volume (MCV) and (**D**) Mean Corpuscular Hemoglobin (MCH) in CC037*ΔF508/ΔF508* (HO) and *wt/wt* (WT) mice at PND28. Data are expressed as mean values ± standard errors of the means (SEM). Data represents the following genotypes and genders: *n*=13 WT and *n*=12 HO mice (number of males: WT=7, HO=7 and number of females: WT=6, HO=5). Statistical significance, determined by Mann-Whitney test non-parametric test two-tailed, is indicated as follows: ****, p<0.0001. **E-F)** H&E stained sections of the bone marrow showing mild-to-marked granulocytic hyperplasia of HO compared to that of WT mice at PND 28. H&E staining is representative of *n*=14 WT and *n*=15 HO mice (number of males: WT=7, HO=8; number of females: WT=7, HO=7). Scale bars=20 μm **G**) The cellularity of sternum and femur granulocytes was scored in bone marrow stained with H&E. Values represent the mean ± standard error of the mean pooled from n=11 mice for each group Statistical significance determined by Mann-Whitney test is indicated as follows: ***, p<0.005; ****, p<0.0001.

The bone marrow of CC037*ΔF508/ΔF508* mice, in both the sternum and femur, showed trilineage hematopoiesis with mild-to-marked increased cellularity of the myeloid compartment, with a predominance of mature granulocytes, compared to that of *wt/wt* mice (**Fig. 5E-G**). Thus, the *ΔF508-Cftr* mutation sustains granulocytic hyperplasia in the bone marrow and an increased percentage of neutrophils in the peripheral blood, supporting the presence of a systemic inflammatory response.

Minimal-to-mild extramedullary hematopoiesis, as evidenced by erythroid, myeloid and megakaryocytic precursors in the red pulp, occurred in the spleens of CC037 *ΔF508/ΔF508* with no major differences compared to *wt/wt* mice (**Fig. S3A, B)**. Histopathological analysis did not reveal obvious differences in the thymus of CC037*ΔF508/ΔF508* as compared to *wt/wt* mice (**Fig. S3C, D**).

### Morphological changes were present in the pancreas, heart, and male reproductive tracts of CC037ΔF508/ΔF508 mice

The pancreas of CC037*ΔF508/ΔF508* mice showed several hypoplastic and under-developed islets, composed of few numbers of small and poorly defined endocrine epithelial cells (**Fig. 6A**), in contrast to well-developed and structured islets in *wt/wt* mice (**Fig. 6B**). Although the total area of pancreas from CC037*ΔF508/ΔF508* and *wt/wt* mice was not significantly different (**Fig. 6C** and **Fig. S3E**), the total area of pancreatic islets was significantly lower in CC037*ΔF508/ΔF508* compared to CC037 wt/wt mice (**Fig. S3F**). In particular, CC037*ΔF508/ΔF508* mice had both a smaller average size of pancreatic islets (**Fig. 6D**), and a lower number of islets compared to *wt/wt* mice (**Fig. 6E**). These differences justify the presence of a smaller relative area of the islets on the entire pancreas in the CC037*ΔF508/ΔF508* compared to the *wt/wt* mice (**Fig. 6F**).

**Fig. 6:**
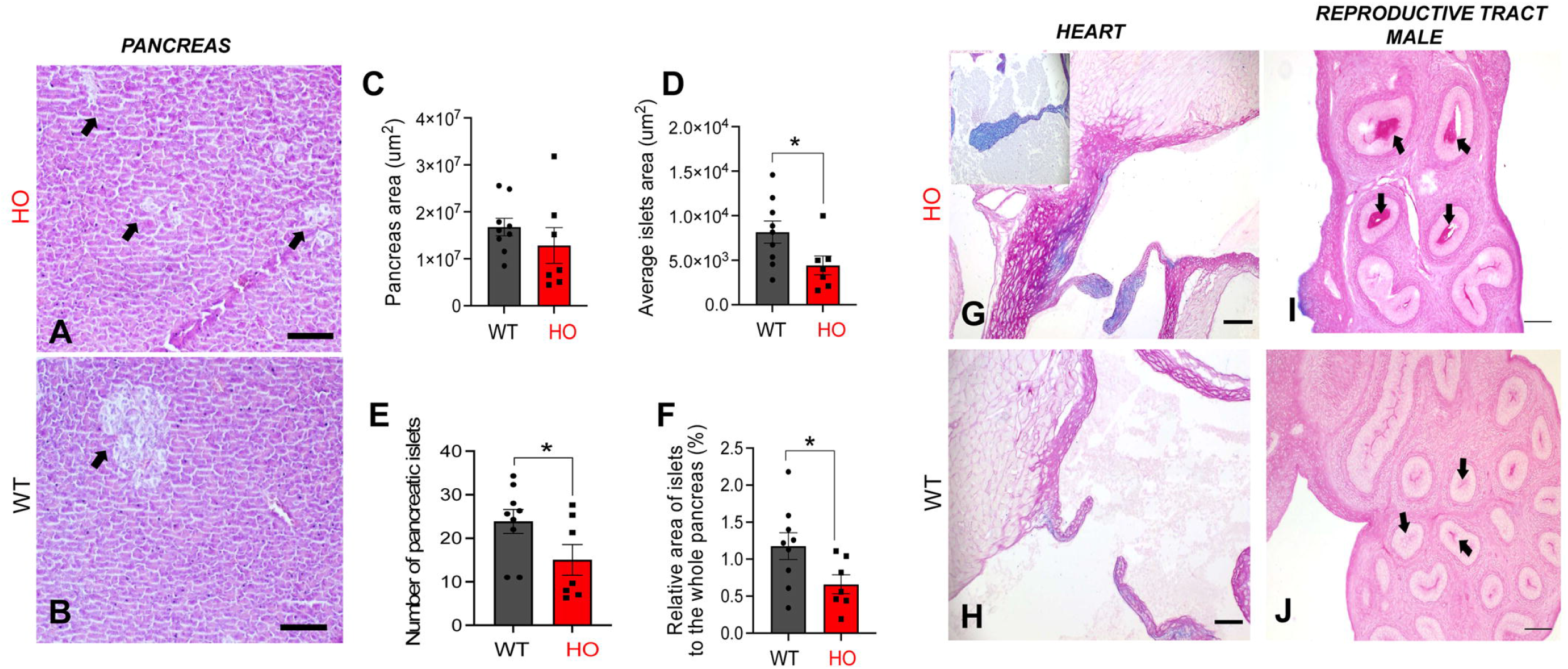
Disease phenotypes in the pancreas, heart and epididymis of CC037 *ΔF508/ΔF508* and CC037 *wt/wt* mice. **A-B)** H&E staining of the pancreas at PND28. Hypoplastic and poor developed pancreatic islets, composed by only few numbers of small and poorly defined endocrine epithelial cells (**A,** arrowheads) in *CC037ΔF508/ΔF508* (HO), compared to well-developed and structured islets in CC037 *wt/wt* (WT) mice (**B,** arrowheads). **C**) Total area of the pancreas, (**D**) the average size of pancreatic islets, (**E**) the number of islets, (**F**) relative area of the islets on the entire pancreas in HO compared to WT mice. Values represent the mean±SEM pooled from n=9 WT and n=7 HO mice (number of males: WT=4, HO=4; number of females: WT=5, HO=3). Statistical significance determined by Mann-Whitney test is indicated as follows: *p < 0.05, **p < 0.01, ***p<0.005. **G**) AB-PAS staining of the hearts at PND28. HO mice showed endocardiosis with left mitral valve thickening (insert in **G**) compared to normal appearance in WT (**H**) mice. **I**) AB-PAS staining of testes showing immature spermatocytes of HO mice compared to round and elongating spermatids of WT (**J**). **K**) AB-PAS staining of epididymis showing abundant PAS-positive material (arrows) in the tubules of HO mice compared to morphologically normal epididymal tubules of WT (**L,** arrows). AB-PAS staining is representative of *n*=14 WT and *n*=15 HO mice (number of males: WT=7, HO=8; number of females: WT=7, HO=7). Scale bars= 250 μm (**A, B**, **G, K, H, L**); 20 μm (**I, J,**).

Endocardiosis was occasionally observed in the hearts of CC037*ΔF508/ΔF508* mice (**Fig. 6G**). Atrioventricular valves, in particular right–sided, showed thickening of the leaflets due to accumulation of ground matrix, rich in sulphated and carboxylated acid mucopolysaccharides and glycoproteins as highlighted by AB-PAS staining. This change was also observed at the level of the aortic and pulmonary valves, although less marked at those sites. In contrast, this change was absent in *wt/wt* mice (**Fig. 6H**).

Testes of CC037*ΔF508/ΔF508* mice were more sexually immature (**Fig. 6I**), with seminiferous tubules showing zygotene/pachytene spermatocytes compared to their age-matched peripubertal *wt/wt* littermates (**Fig. 6J**), with round and elongating spermatocytes and occasionally spermatids.

In CC037*ΔF508/ΔF508* male mice, the lumen of epididymis was characterized by abundant PAS-positive material (**Fig. 6K)**. compared to lument of epididymis of *wt/wt* mice (**Fig. 6L)**. Thus, *ΔF508-Cftr* mutation could determines several early pathological features that may lead to severe abnormalities in later developmental stages.

## Discussion

This is the first report utilizing a highly genetically diverse CC mouse population to produce genetically modified mice with *ΔF508-Cftr* mutation, demonstrating that they can develop different pathological phenotypes never described in previously-characterized mouse models of CF (**Fig. 7**). While CC lines have been used to model a wide range of biomedically relevant traits ^18,19^, they have never been used to obtain a specific gene mutation. Generally, these CC lines have high variations in breeding productivity and fertility ^25^, which may be further compromised by insertion of *ΔF508-Cftr* mutation. We have mitigated these limitations by selecting the CC006 and CC037 lines from a panel of 39 CC lines based on their fertility and reproductive fitness (http://csbio.unc.edu/CCstatus). In addition, key relevant traits for selection were the high number of different founder strains to ensure genetic variability, low heterozygosity to assure genetic stability and the absence of spontaneous pathological phenotypes to exclude confounding factors.

**Figure 7:**
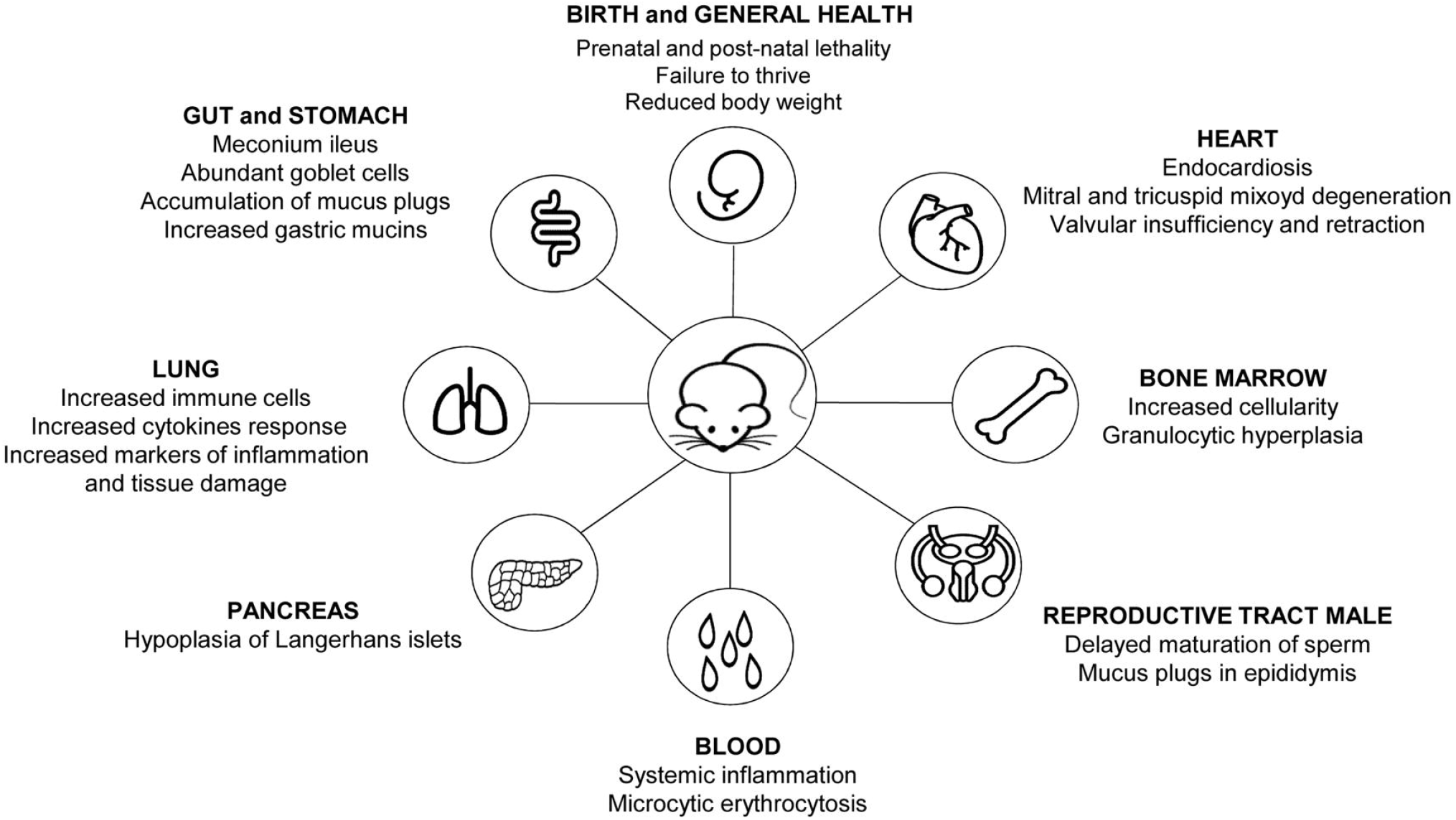
Schematic view of major pathological changes in *CC037ΔF508/ΔF508* in comparison to *wt/wt* mice.

Susceptibility to infection by pathogens, including *P. aeruginosa*, were criteria to reproduce the CF disease phenotype ^20,21,26,27^. The *ΔF508-Cftr* mutation was introduced into the CC006 and CC037 lines by backcrossing with the existing C57BL/6J Cftr^tm1Kth^ mouse line ^24^. Overall, the fertility in heterozygotes CC006 murine line is poorer compared to the CC037 line. Accordingly, sequencing data showed that the CC006 line carries both the CAST/EiJ and PWK/PhJ haplotypes, which have been previously shown to contribute to breeding difficulties and poor fertility ^28^. Breeding *CC006ΔF508/wt* heterozygotes does not produce *CC006ΔF508/ΔF508* homozygotes, while breeding CC037*ΔF508/wt* heterozygotes produces an average of 13% CC037*ΔF508/ΔF508* homozygotes. The percentage of *ΔF508/ΔF508* mice in the CC037 line is significantly different from expected and from that in the C57BL/6J Cftr^tm1Kth^ mice used for backcrossing ^24^. For the first time, our results in CC lines carrier of *Cftr-ΔF508* mutation showed an unexpectedly increased risk of prenatal lethality for *ΔF508* homozygotes, which raises several key questions regarding human pathogenesis. This prenatal lethality suggests a notably underestimated impact of host genetic variations together with *Cftr* mutation on implantation or embryo development. The CFTR mRNA transcript has been found in human fetuses as early as ten weeks gestation ^29^ and during early human embryo development where the CFTR functions in apical membranes of the human blastocyst contribute to the early development of human embryo ^30^. Mortality extended into the postnatal period, with CC037*ΔF508/ΔF508* homozygotes dying by eight weeks of age. This limited the time span for phenotypic analyses to PND28, and we elected to carry out these analyses in the naïve state, i.e., without confounding effect due to the early introduction of osmotic laxative. The severe lethality and failure to thrive of *CC037ΔF508/ΔF508* mice was more severe than that previously observed with *ΔF508/ΔF508* genotypes in Cftr^tm/Eur^ (10%) ^31^ and *Cftr*^tm1Kth^ mice (60%) ^24^ but similar to that observed in *Cftr*^tm2Cam^ (95%) transgenic mice ^32^.

To understand genomic differences between CC037 and CC006 murine strains with *ΔF508-Cftr* mutation established in this study and previously reported CF mice, we performed sequence analysis. CC037 and CC006 transgenic murine strains have the NOD/ShiLtJ and NZO/H1LtJ founder haplotypes for the *Cftr* locus, respectively, which makes these lines substantially different from those previously reported with the *Cftr* mutation in C57BL/6J. Our *ΔF508* transgenic CC lines have residual heterozygosity below 10%, which is in the range recommended for the distribution of CC lines and valuable for genetic mapping, validation experiments, and disease modeling ^33^. However, because the CC037*ΔF508/ΔF508* line was not fully inbred and residual heterozygosity may also influence experimental replication, our phenotypic analyses were expanded to multiple mice from different generations and different breeding pairs.

One of the key yet unanswered questions in CF is what comes first between lung infection and inflammation. Differently from previous CF mouse models, CC037*ΔF508/ΔF508* mice show early neutrophil-dominated inflammation recapitulating features that likely initiate lung disease in pwCF. Inflammation is a key participant in the lung destruction observed in pwCF; however, in humans, the temporal and causal relationships between inflammation and infection remain unclear. There are conflicting data from the bronchoalveolar lavage fluid of infants and children with CF regarding the presence of inflammation and infection early in CF ^34–36^. Animal models of CF have yielded data both supporting and opposing inherent inflammation. The porcine model of CF showed no evidence of inflammation or infection a few hours after birth, while later, porcine lungs contained multiple bacterial species and exhibited airway inflammation ^13^. Additionally, CF ferrets rapidly develop lung disease characterized by bacterial colonization ^10^, although lung disease persists after bacterial eradication. While data from previous mouse models do not address this question, our data in the CC037 murine background indicated that the *ΔF508-Cftr* mutation led to an early intrinsic pro-inflammatory state in developing CF airways.

A key component of CF muco-obstructive lung disease is the presence of airway mucus accumulation. We found sporadic mucus plaques in the respiratory tracts including nasal and paranasal sinuses, trachea, and lung of *CC037ΔF508/ΔF508* mice. Thus, we can not conclude that mucus plugging is the main determinant of lung inflammation in this model. Similar to previous CF mouse models, CC037*ΔF508/ΔF508* mice showed significant mucus impaction in the gut, with necrotic and fecal material deposition in the small intestine. This severe intestinal obstruction significantly impacts survival and can be the cause of systemic inflammation with the spreading of inflammatory stimuli to distant organs. In this regard, the hematological analysis of CC037*ΔF508/ΔF508* mice had a significantly higher neutrophil percentage and lower lymphocytic counts than *wt/wt* mice, indicating ongoing neutrophil-driven systemic inflammation in young CF mice.

The mortality of CF mice was attributed to severe intestinal obstruction ^24^. Our phenotypic characterization shows consistent postnatal intestinal complications in CC037*ΔF508/ΔF508* mice as in previous models. In addition, we describe an additional prevalent and severe phenotype in the hearts of CC037*ΔF508/ΔF508* mice that may play a role in mortality. The *ΔF508* mutation in CC037 mice was associated with endocardiosis with valve remodeling that may lead to cardiovascular disturbances, potentially causing arrhythmia-related syncope or sudden death. With the prominent focus on pulmonary disturbances in humans and gut pathology in animal models, the impact of cardiovascular disturbances in pwCF has been underestimated. The majority of studies in humans with CF have focused on altered right ventricle function as a consequence of increased pulmonary pressures in individuals with lung disease ^37,38^. Some studies in mice have shown that left ventricular and aortic functions are affected by *Cftr* loss in the absence of lung disease ^39^, while the cardiovascular disease has not been reported in large CF animal models. Overall, the role of CFTR in the heart remains to be clarified, but our findings provide a compelling reason to expand studies on the cardiovascular function of pwCF early in life.

Exocrine pancreatic insufficiency is a major condition of pwCF while in small animal models of CF, including mice and rats, only those carrying null CFTR mutations (CFTR^tm1UNC^ and CFTR^tm1CAM^) exhibit mild pancreatic changes, with variable penetrance, and they do not exhibit dramatic alterations in morphology ^40–42^. The *ΔF508* mutation in C57Bl/6 mice (CFTR^tm2CAM^, CFTR^tm1EUR^, CFTR^tm1KTH^, and CFTR^tm1G551D^) does not cause pancreatic abnormalities. In our study, the *ΔF508* CFTR mutation in the CC037 background showed both a lower average size and number of pancreatic islets compared to *wt/wt* mice, despite having pancreases of comparable size. Although these changes are milder than those observed in the pig and ferret models ^10,43^, they are still significantly different from those in previous *ΔF508* mouse models.

Pathological liver and gallbladder changes have been documented in pwCF. In particular, focal biliary cirrhosis is the second most common cause of mortality in pwCF. Similar to previous mouse models of CF, no remarkable liver and gallbladder abnormalities in CC037 *ΔF508/ΔF508* mice were documented in our study.

Renal involvement in CF may be associated with comorbidities and treatment rather than a direct effect of CFTR mutation. The most well-known cause of renal injury in pwCF is iatrogenesis, which is related to the intake of high doses of aminoglycosides during pulmonary exacerbation ^44^.The kidneys of naïve CC037 *ΔF508/ΔF508* mice were unaffected indicating that CFTR mutation may not contribute *per se* to renal disease.

Human bone marrow produces hematopoietic cells that contribute to the inflammatory response ^45^. Although CF is characterized by both pulmonary and systemic inflammation, as well as by defective lung repair, there have been no studies or evidence supporting the hypothesis that bone marrow function in pwCF is altered. However, studies in CF mice showed that bone marrow transplant from *Cftr-wt/wt* donors significantly improves survival and reduces inflammation in *ΔF508-Cftr* mice providing immune support and rationale for cell-based therapy ^46^. In our study, the bone marrow of CC037*ΔF508/ΔF508* mice showed mild-to-marked granulocytic hyperplasia while *wt/wt* mice did not. Neutrophil-driven systemic inflammation in CC037*ΔF508/ΔF508* mice and microcytic erythrocytosis related to *ΔF508-Cftr* mutation have been described here for the first time. Hematological analysis showed that CC037*ΔF508/ΔF508* mice had a high number of red blood cells with a reduced size and a reduced content of hemoglobin compared to *wt/wt* mice. The erythrocytosis could be caused by different factors including defects in bone marrow, cardiac disease or caused by other factors including hypoxia and breathing difficulty ^47^. In addition, other conditions that influence a reduced MCV and are possibly related to our mouse model are iron deficiency but also chronic disease, inflammation, and infections ^48^.

Male infertility in pwCF is mainly caused by congenital absence of the vas deferens resulting in azoospermia and/or poor sperm quality ^49^. Previous mouse models failed to recapitulate this infertility phenotype ^40,42,50,51^. In piglets, the vas deferens appeared to be intact ^52^, but it was absent or degenerate in knockout ferrets ^10^. CC037*ΔF508/ΔF508* male mice showed pathological epididymal tubules with a greatly reduced concentration of mature sperm supporting possible infertility at later development.

Taken together, our model focused on *Cftr*-ΔF508 mutation in a diverse genetic profile covers novel phenotypes previously undocumented in small animal CF models, including prenatal mortality, and abnormalities in lung, blood, pancreas, bone marrow, heart, and male reproductive system. Notably, research efforts focusing on these extra-pulmonary organs have become topical in the era where highly effective CFTR modulator therapies have significantly ameliorated pulmonary manifestations. Our study strongly suggests that genetic background can influence the development of currently unidentified CFTR-dependent disease phenotypes. Development of these novel CF mouse models is expected to facilitate progress toward a more detailed understanding of CF pathogenesis and support the identification of diverse disease manifestations in the human population.

## Materials and Methods

### Animals and ethics statement

Animal studies adhered to the Italian Ministry of Health guidelines for the use and care of experimental animals (IACUC 739, 935, and 1043). CC037 and CC006 mice were obtained from the UNC Systems Genetics Core Facility at UNC at Chapel Hill. C57BL/6 *ΔF508/wt* (Cftr^tm1Kth^) mice ^24^ were obtained from Case Western Reserve University.

The *ΔF508-Cftr* mutation was backcrossed into CC006 and CC037 mice for five generations using marker-assisted accelerated backcrossing (MAX-BAX®) ^53^ as shown in **Fig. S1.** Experimental cohorts of mice were produced by intercrossing heterozygous (*ΔF508/wt*) mice to obtain mice of *wt/wt*, *ΔF508/wt*, and *ΔF508/ΔF508* genotypes. Breeding pairs had no mother and father or grandparents in common. At birth, all pups were toe-clipped for identification and genotyping was performed at death and 7-10 PND.

### Genomic analysis

*CC006ΔF508/wt* and *CC037ΔF508/wt* were selected for WGS. One hundred nanograms of DNA obtained from tails were used for library preparation with a TruSeq DNA Nano kit (Illumina). Libraries were sequenced on the NovaSeq 6000 platform (2 x 150 bp) at CeGaT GmbH (Tübingen, Germany). In total, 92.82% of the reads had a QC value > 30. Demultiplexing of the sequencing reads was performed with Illumina bcl2fastq (version 2.20). Trimming was performed with BBMap bbduk v38.68 [BBMap - Bushnell B. - sourceforge.net/projects/bbmap/] (with parameters k=23 mink=11 rcomp=t ktrim=f kmask=N qtrim=rl trimq=5 forcetrimleft=5 forcetrimright2=0 overwrite=true) to remove adapter sequences and low-quality bases from the reads. Clean reads in FastQ format were aligned to the reference mouse genome mm10 (C57BL/6J) by Burrows-Wheeler Aligner (bwa-mem2 v2.2.1) ^54^. Duplicate reads were highlighted with picard MarkDuplicates v2.24.2 [http://picard.sourceforge.net/] after sorting and indexing of bam alignment files performed using samtools v1.11 ^55^.

SNP and small indel variants were detected according to the GATK4 (O’Connor and Auwera 2020) best practice guidelines (https://gatk.broadinstitute.org/hc/en-us/articles/360035535932-Germline-short-variant-discovery-SNPs-Indels-). Variant flags were assigned using SnpSift utilities (4.1g) (Cingolani et al. 2012) to identify (a) fixed differences from the reference that are present across all CC strains, (b) variants previously reported (Keane et al. 2011). Finally, variant annotation was performed by snpEFF (4.1g) ^56^ and the highest impact annotation for any variant was retained.

To reconstruct the haplotype contributions from the eight founders, we used R/qtl2 v0.24 ^57^ in the R environment (v. 3.6.3) as described in Supplementary information.

### Data and Code Availability

WGS data generated in this study have been deposited in SRA, accession number PRJNA915352. All codes associated with this manuscript have been uploaded to GitHub https://github.com/volpesofi/CCmice_scripts and are available in Zenodo at the following doi 10.5281/zenodo.7521696.

### Flow cytometric and other analyses of the lung

Lung single-cell suspensions were obtained by mashing the lungs through a 70-μM cell strainer in RPMI+5% FBS. All the procedures were previously described ^58^ and reported in the online data supplement. Mucin agarose western blot was performed on reduced and unreduced BAL samples as previously described ^59^. Proteins were measured using the Bradford assay. MPO and neutrophil elastase were measured in the lung supernatants by ELISA (R&D System). Cytokine/chemokine levels were measured in supernatant of lung homogenate by Bioplex Assay (Bio-rad).

### Blood analyses

Blood was collected from the retro-orbital plexus of anesthetized animals using Na-heparin coated capillaries (Hirschmann Laborgera□te GmbH) and vials (Microvette® 200 K3 EDTA, 20.1288 Sarstedt). Hematologic parameters were evaluated using an automated cell counter (ProCyte Dx, IDEXX Laboratories, USA).

### Histology

Lung, nasal cavity, trachea, stomach, gut, liver, gallbladder, pancreas, kidneys, heart and reproductive tracts were collected and fixed in Carnoy solution. The spleen, thymus, sternum, and femur-tibia were collected and fixed in 10% neutral buffered formalin. The femur-tibia, sternum and nasal cavity were decalcified using a solution of 37% hydrochloric acid and 98% formic acid for 3 hours. Tissues were embedded in paraffin, cut, and stained with H&E and AB-PAS. Microscopic lesions were classified on a scale of 1 to 5 as minimal (1), mild (2), moderate (3), marked (4), or severe (5); minimal referred to the least extent discernible, and severe referred to the greatest extent possible. The total area of the pancreas and the islet’s area were scored with Aperio ImageScope Software.

### Statistics

Statistical analyses were performed with GraphPad Prism using Mantel-Cox test for survival curve and Mann-Whitney two-tailed comparison test for the other readouts. Outlier data, identified by Grubbs’ test, were excluded from the analysis.

## LIST OF SUPPLEMENTARY MATERIALS

**Fig. S1. Backcross strategy to generate CC037*ΔF508/ΔF508* and CC006*ΔF508/ΔF508* mice.**

**Fig. S2. Respiratory disease phenotype of CC037 *ΔF508/ΔF508* and *wt/wt* mice.**

**Fig. S3. Spleen, thymus and pancreas phenotype in CC037 *ΔF508/ΔF508* in comparison with CC037 *wt/wt* mice.**

**Table S1. Criteria for selection of CC lines from database (http://csbio.unc.edu/CCstatus) and other publications.**

**Table S2. Progeny obtained from the cross of CC006 and CC037 lines heterozygous for ΔF508.**

**Table S3. Mortality of the CC037 and CC006 offsprings from *ΔF508-Cftr* heterozygous intercrosses.**

**Table S4. Cytokines and chemokines in lungs of CC037*ΔF508/ΔF508* and *wt/wt* mice at PND28.**

**Table S5. Haematological analysis of CC037*ΔF508/ΔF508* and *wt/wt* mice at PND28.**

## Acknowledgments

We thank R. Norata, F. Colombi, F. Saliu, G. Rizzo, G. Larosa and M. D’aurora for technical support and I. De Fino and M. Ferris for useful discussion and critical reading of the manuscript. We thank OSR Mouse Clinic, OSR animal biochemistry for hematological analysis and FRACTAL for FACS analysis.

## Author contributions

B. Sipione performed experiments, analyzed data, performed the statistical analyses, prepared the figures and edited the paper. N.I. Lorè performed experiments, analyzed data, provided funding and edited the paper. G. Rossi and F. Sanvito C. performed experiments, analyzed data, provided conceptual advice and wrote paper. A. Neri performed experiments, analyzed data, performed the statistical analyses, prepared the figures and edited the paper. F. Gianferro performed experiments, analyzed data, performed the statistical analyses, prepared the figures and edited the paper.A. S. Tascini performed experiments, analyzed data and wrote paper. A. Livraghi-Butrico A. performed experiments, analyzed data, provided conceptual advice and wrote paper. C. Cigana performed experiments, analyzed data, provided conceptual advice and edited the manuscript. Bragonzi designed and coordinated the study, analyzed data, provided funding, and wrote the paper.

