## Supplementary File Methods for "*ΔF508-Cftr* mutation in genetically diverse Collaborative Cross mice yields novel disease-relevant phenotypes for cystic fibrosis"

Supplementary Methods, Tables and Figure legends

**SUPPLEMENTARY METHODS**

**Animals**. Mice were maintained in the pathogen-free facility at San Raffaele Scientific Institute (Milano, Italia) where 3-5 mice per cage were housed. Mice were maintained in sterile ventilated cages. Mice were fed with standard rodent autoclaved chow (VRFI, Special Diets Services, UK) and autoclaved tap water. Fluorescent lights were cycled 12h on, 12h off, and ambient temperature (23±1°C) and relative humidity (40-60%) were regulated.

The *Cftr*-ΔF508 mutation was backcrossed into CC006 and CC037 mice for five generations using marker-assisted accelerated backcrossing (MAX-BAX®) according with the breeding scheme shown in **Fig. S1.** At each generation, we selected progeny with the highest percentage of CC background as breeders for the next generation and to five generations to produce the congenic strains CC006 *ΔF508/wt* and CC037 *ΔF508/wt.*

The primers used to amplify the endogenous and mutant *Cftr* loci were TTC AAG CCC AAG CTT TCG CGA G (mutant), CTC CCT TCT TCT AGT CAC AAC CG (common), and CAT CTT GAT AGA GCC ACG GTG C (wild type). The mice studied were age-matched littermates of both sexes.

**Genomic analysis.** Quality check of raw sequencing data was performed with FastQC v0.11.9 <http://www.bioinformatics.babraham.ac.uk/projects/fastqc/>. SNP and small indel variants call was performed with gatk HaplotypeCaller (v. 4.1.9). The mm10 resource bundle for base recalibration with gatk4 was built as described at https://github.com/igordot/genomics/blob/master/workflows/gatk-mouse-mm10.md. Specifically, mouse indels were downloaded from the Sanger Mouse Genetics Programme (5/2015 release), while mouse SNPs were downloaded from NCBI (organism: mouse_10090) and converted from GRCm38 to mm10 format. After variants calling, the single sample vcfs were merged with the vcf-merge utility of vcftools (0.1.16). The percentage of heterozygosity in the autosomes of sequenced mice was assessed with bcftools stats (v 1.9) and computed as the ratio between heterozygous SNPs (nHet) and the total number of SNPs (nHet + nRefHom + nNonRefHom).

vcftools utility tools were also used to filter vcf on MegaMuga GigaMuga positions (--positions). Information on the genomics location of the MegaMuga GigaMuga genotypes was retrieved from the MMnGM_pmap*.csv files downloaded from figshare (https://doi.org/10.6084/m9.figshare.5404762.v2).

**Haplotype reconstruction.** Haplotype contributions from the eight founders were calculated with R/qtl2 v0.24 (Broman KW et al. 2019) in the R environment (v. 3.6.3)- VCF files were firstly filtered on the MegaMUGA and GigaMUGA (combined) SNP genomics location used for Collaborative Cross genotyping using vcftools, then imported in R with data.table (v 1.14.0), prepared with qtl2convert (v0.24) and merged with the original genome sequences of the CC lines (Srivastava et al. 2017) retrieved from Zenodo (https://doi.org/10.5281/zenodo.377036). Founders’ genotypes for use with R/qtl2 were retrieved from figshare (https://doi.org/10.6084/m9.figshare.5404762.v2). For each sequenced individual, the observed fraction of the autosomes assigned to each of the 36 possible diplotype states were computed with Rqtl2/calc_genoprob function. We then reduced genotype probabilities to the allele probabilities of each founder haplotype using Rqtl2/genoprob_to_alleleprob function. A haplotype mosaic plot for each sample was created with ggplot2 (v. 3.3) from the allele probability data-frame rounded to 0, 0.5 or 1. The obtained haplotype mosaic plot was compared with the original CC line, CC037 and CC006 respectively, and the small differences detected were manually inspected. The density of heterozygous variants was used to check the reliability of haplotype reconstruction by Rqtl2

**Lung flow cytometric analysis.** After centrifugation of the mashed lung cell suspension, the surnatants were recovered and stored at -80°C for the cytokine analysis. Lung cells (3x10^6^ cells) were incubated with blocking buffer (5% rat serum and 95% culture supernatant of 24G2 anti-FcR mAb-producing hybridoma cells) for 10 min at 4°C. Then, cells were stained for 20 min at 4°C in the dark with different combinations of the following antibodies (BD Biosciences): CD45- PacBlue (clone30-F11), Gr-1- FITC (clone RB6-8C5), CD11b- APC (clone M1/70), CD11c- PeCy7 (clone N418), B220- PerCP (clone RA3-6B2) and I-A/I-E-PE (clone M5/114.15.2).

After the staining, cells were fixed (paraformaldehyde 2% in PBS for 10 min) and resuspended in 500 µl of PBS. Acquisition and analyses were performed using the analyzer Cytoflex S (Beckman Coulter) and FlowJo Software. CompBeads (BD Biosciences #552845 Anti-Rat and Anti-Hamster Ig ĸ /Negative Control Compensation Particles Set) were used to prepare singlets and negative control. One drop of CompBeads and one drop of Negative control were added in each tube, stained for 20 min at 4°C in the dark with different antibodies and resuspended in 500 µl of saline solution.

**SUPPLEMENTARY FIGURE**

**Fig. S1.** **Backcross strategy to generate CC037*ΔF508/ΔF508* and CC006*ΔF508/ΔF508* mice.** The *Cftr*-ΔF508 mutation of C57BL/6J *ΔF508/wt* (Cftr^tm1Kth^) mice was backcrossed (BC) into CC06 and CC037 murine lines for five generations using Marker-Assisted Accelerated Backcrossing (MAX- BAX®). Experimental cohorts of mice were produced by intercrossing heterozygous (*ΔF508/wt*) mice to obtain mice of three different genotypes with respect to murine Cftr (*wt/wt*, *ΔF508/wt* and *ΔF508/ΔF508*).

**Fig. S2. Respiratory disease phenotype of CC037*ΔF508/ΔF508* and *wt/wt* mice.**

**A**) Representative pictures of the respiratory tract at PND28. **B-E**) AB-PAS staining at PND28. Turbinates of CC037*ΔF508/ΔF508* (HO) (**B,** **C**) showed increased mucus secretion (arrows) compared to CC037*wt/wt* (WT) mice (**F, G**). The trachea, sampled immediately below the larynx, showed an increased amount of goblet cells (arrows in **D**) and hyperplastic/hypersecretory appearance of the peri-laryngeal glands (arrowheads) in HO mice compared to WT (**H**). Lung of HO mice was highly heterogeneous showing lobes with focal accumulation of mucus adhering to the airway and marked concentration of goblet cells (open-arrows) (**E**). The absence of pathology with no mucus plugs and a thin monolayer of epithelial cells was observed in the lung of WT mice (**I**). Scale bars= 500 µm (B-F); 50 µm (C,G); 250 µm (D,E,H,I). AB-PAS stain is representative of *n*=14 WT *and 15* HO mice (n male: WT=7, HO=8*;* n female WT=7*,* HO=7*)*.

**Fig. S3. Spleen, thymus and pancreas in CC037*ΔF508/ΔF508* in comparison with *wt/wt* mice. A-D)** H&E stained sections of the spleen (**A, B**) and thymus (**C,D**) shows similar histologic pattern between CC037*ΔF508/ΔF508* and CC037*wt/wt* mice. Scale bar 50 $\mu$m (**A, B**); 100 $\mu$m (**C, D**).

H&E stain is representative n=14 wt/wt and 15 ΔF508/ΔF508 mice (n male: wt/wt=7, ΔF508/ΔF508=8; n female wt/wt=7, ΔF508/ΔF508=7).

**E**) Total area of the pancreas normalized for mice weight; **F**) Area of the pancreatic islets in CC037 *ΔF508/ΔF508* compared to *wt/wt* mice.

**Table S1. Criteria for selection of CC lines from database (**[**http://csbio.unc.edu/CCstatus**](http://csbio.unc.edu/CCstatus)**) and other publications.**

**Table S2. Progeny obtained from the cross of CC006 and CC037 lines heterozygous for *Cftr-ΔF508*.**
