## Supplementary Table for "*ΔF508-Cftr* mutation in genetically diverse Collaborative Cross mice yields novel disease-relevant phenotypes for cystic fibrosis"

**Table S1. Criteria for selection of CC lines from database (**[**http://csbio.unc.edu/CCstatus**](http://csbio.unc.edu/CCstatus)**) and publications.**

| **CC line ID** | **Nr of founder strains** | **% heterozygosity in MRCA^§^** | **Breeding well**  **in multiple female cages** | **Litter Size** | **Spontaneous Disease** | ***P. aeruginosa***  **response*** | **Other pathogens response^#^** | **Score** |
| --- | --- | --- | --- | --- | --- | --- | --- | --- |
| **CC043** | 8 | 4.7 | yes | 3,5 |  | susceptible |  | 6 |
| **CC072** | 6 | 0.8 | yes | 4,1 | Pups from the same litter are born at different sizes | susceptible |  | 5 |
| **IL-1061** | - | - | - | - | - | susceptible |  | 2 |
| **CC076** | 8 | - | - | - | - | susceptible |  | 4 |
| **CC051** | 6 | 6,9 | - | 6,6 | Males are prone to rectal prolapse. | susceptible | *Klebsiella pneumonia*  *Aspergillus fumigatu*s | 5 |
| **IL-4438** | 6 | - | - | - | - | susceptible |  | 3 |
| **IL-1513** | - | - | - | - | - | susceptible |  | 2 |
| **IL-4052** | - | - | - | - | - | susceptible |  | 2 |
| **CC037** | 6 | 1,1 | yes | 5,8 | - | susceptible | *Klebsiella pneumonia*  *Aspergillus fumigatu*s | 7 |
| **CC019** | 6 | 9 | yes | 5,4 | Jumpy & aggressive. High incidence of hydrocephalus | susceptible |  | *5* |
| **IL-611** | - | - | - | - | - | susceptible |  | *2* |
| **CC016** | 7 | 3,7 | yes | 4,1 | - | intermediate |  | *5* |
| **CC059** | 6 | 10,3 | - | 5,6 | - | intermediate |  | *3* |
| **CC028** | 8 | 2,1 | yes | 4,3 | Obese when adults and stop reproducing.  Jumpy when young. | intermediate |  | *4* |
| **IL-111** | - | - | - | - | - | intermediate |  | *1* |
| **IL-4457** | - | - | - | - | - | intermediate |  | *1* |
| **CC013** | 7 | 0,5 | yes | 5,4 | - | intermediate |  | *5* |
| **CC033** | 8 | 9,3 | no | 3,7 | - | intermediate |  | *4* |
| **CC010** | 8 | 3,4 | yes | 4 | - | intermediate |  | *5* |
| **CC041** | 6 | 9,5 | - | 5,8 | - | intermediate |  | *4* |
| **CC018** | 8 | 9,6 | yes | 3,5 | - | intermediate |  | *5* |
| **IL-519** | - | - | - | - | - | intermediate |  | *1* |
| **CC011** | 8 | 5,9 | yes | 4,7 | Prone to colitis and rectal prolapses. | intermediate |  | *4* |
| **IL-2131** | - | - | - | - | - | intermediate |  | *1* |
| **CC012** | 8 | 9,3 | yes | 3,6 | - | intermediate |  | *5* |
| **IL-2156** | - | - | - | - | - | intermediate |  | *1* |
| **CC005** | 8 | 1,1 | yes | 3,3 | Males are prone to prepucial gland abscesses. | intermediate |  | *4* |
| **CC002** | 8 | 10,2 | yes | 3,9 | - | intermediate |  | *4* |
| **CC006** | 8 | 8,3 | yes | 3,8 | - | resistant |  | 6 |
| **CC017** | 8 | 7,4 | yes | 4,1 | Males can be very aggressive. | resistant |  | 5 |
| **CC024** | 8 | 3,6 | yes | 4,1 | Jumpy | resistant |  | 5 |
| **CC036** | 8 | 9 | no | 3,1 | - | resistant |  | 5 |
| **CC039** | 8 | 10,3 | yes | 3 | Jumpy | resistant |  | 4 |
| **CC040** | 8 | 11,4 | - | 4 | - | resistant |  | 4 |
| **CC042** | 7 | 6,3 | no | 5,3 | Prone to dermatitis | resistant |  | 4 |
| **CC055** | 8 | 10,6 | - | 3 | - | resistant |  | 4 |
| **CC061** | 8 | 5,1 | yes | 3,8 | Occasional muscular dystrophy and a split pubis | resistant |  | 4 |
| **IL-3438** | - | - | - | - | - | resistant |  | 2 |
| **IL-670** | - | - | - | - | - | resistant |  | 2 |

In gray, positive criteria for selection of CC mice to backcross with C57BL/6J (Cftr^tm1Kth^) mice.

In green and red, CC mice selected for backcrossing with final score.

**^§^** According to data derived from dense genotyping of the Most Recent Common Ancestor (MRCA) ^1^.

* Response to *P. aeruginosa* infection (^2,3^)

**^#^** Response to other infection as assessed for *Klebsiella pneumonia* ^4^ and *Aspergillus fumigatu*s ^5^

**Table S2. Progeny obtained from the cross of CC037 and CC006 lines heterozygous for *ΔF508*.**

|  |  | **Genotype observed** | | |
| --- | --- | --- | --- | --- |
| **Cross** | **Nr litters** | ***ΔF508/ΔF508*** | ***ΔF508/wt*** | ***wt/wt*** |
| CC037  *ΔF508/wt* x *ΔF508/wt* | 5,3 | n=369  (13%) | n=1655  (58%) | n=816  (29%) |
|  |  | **Genotype expected** | | |
|  |  | n=710  (25%) | n=1420  (50%) | n=710  (25%) |

|  |  | **Genotype observed** | | |
| --- | --- | --- | --- | --- |
| **Cross** | **Nr litters** | ***ΔF508/ΔF508*** | ***ΔF508/wt*** | ***wt/wt*** |
| CC006  *ΔF508/wt* x *ΔF508/wt* | 3,5 | n=0  (0%) | n=174  (64%) | n=99  (36%) |
|  |  | **Genotype expected** | | |
|  |  | n=68.25  (25%) | n=136.5  (50%) | n=68.25  (25%) |

**Table S3. Mortality of the CC037 and CC006 offsprings from *ΔF508-Cftr* heterozygous intercrosses.**

|  |  | **Mortality** | | | **Survival** |
| --- | --- | --- | --- | --- | --- |
| **Cross** | **Genotype** | **< 5 PND** | **7-21 PND** | **21-28 PND** |  |
| CC037  *ΔF508/wt* x *ΔF508/wt* | ***ΔF508/ΔF508*** | n=96  (29%) | n=58  (17%) | n=109  (33%) | n=69  (21%) |
|  | ***ΔF508/wt*** | n=78  (6%) | n=12  (1%) | n=13  (1%) | N=1268  (92%) |
|  | ***wt/wt*** | n=23  (4%) | n=2  (0,3%) | n=0  (0%) | n=566  (95,7%) |

|  |  | **Mortality** | | | **Survival** |
| --- | --- | --- | --- | --- | --- |
| **Cross** | **Genotype** | **< 5 PND** | **7-21 PND** | **21-28 PND** |  |
| CC006  *ΔF508/wt* x *ΔF508/wt* | ***ΔF508/wt*** | n=5  (9%) | n=0  (0%) | n=0  (0%) | n=49  (91%) |
|  | ***wt/wt*** | n=1  (4%) | n=0  (0%) | n=0  (0%) | n=27  (96%) |

**Table S4. Cytokines and chemokines in lungs of CC037*ΔF508*/*ΔF508* and *wt/wt* mice at PND28.**

| **Cytokine / Chemokine** | **CC037*wt/wt*** | **CC037 *ΔF508*/*ΔF508*** |
| --- | --- | --- |
| IL-1α | 353.93±26.98 | 327.69±37.01 |
| IL-1β | ND | ND |
| IL-2 | 1127.46±50.16 | 1141.45±65.40 |
| IL-3 | ND | ND |
| IL-4 | ND | ND |
| IL-5 | 214.04±18.32 | 315.25±16.85 |
| IL-6 | ND | ND |
| IL-9 | 1482.18±96.96 | 1235.7±111.36 |
| IL-10 | ND | ND |
| IL-12p40 | 1584.23±175.3 | ND |
| IL-12p70 | ND | ND |
| IL-13 | 6153.17±291.48 | 6475.61±435.58 |
| IL-17A | 500.01±20.7 | 617.6±15.6*** |
| Eotaxin | 29236.4±1893.59 | 27883.73±218.67 |
| G-CSF | ND | ND |
| GM-CSF | 1602.85±81.32 | 1537.98±99.49 |
| IFN-γ | 968.86±58.05 | 812.33±75.5 |
| KC | 1719.39±58.44 | 3966.56±750.57** |
| MCP-1 | ND | ND |
| MIP-1α | 70.41±3 | 95.82±8.3** |
| MIP-1β | ND | ND |
| RANTES | 5183.9±425.3 | 4658.44±486.84 |
| TNF-α | 903.53±44.9 | ND |

Lungs of CC037*ΔF508*/*ΔF508* and *wt/wt* mice were collected at 28 PND and physically dissociated as described in material and methods. Pg of cytokines and chemokines normalized for lung weight (g) measured by Bio-Plex assay in lung supernatants. Data are expressed as mean values ± standard errors of the means (SEM). Data represents the following genotypes and genders: *n*=13 *wt/wt* and *n*=12 *ΔF508/ΔF508* mice including number of males: *wt/wt* =7, *ΔF508/ΔF508*=7 and number of females: *wt/wt*=6*, ΔF508/ΔF508*=5. Statistical significance, determined by Mann-Whitney test non parametric test two-tailed, is indicated as follows: *, p<0.05; **, p<0.01; ***, p<0.001. ND, non-detectable

**Table S5. Haematological analysis of** **CC037*ΔF508*/*ΔF508* and *wt/wt* mice at PND28.**

| **Blood count parameters** | **CC037*wt/wt*** | **CC037 *ΔF508*/*ΔF508*** |
| --- | --- | --- |
| RBC (M/µl) | 9.53±0.11 | 10.69±0.16**** |
| HGB (g/dl) | 12.88±0.15 | 13.48±0.19 |
| HCT (%) | 45.13±0.76 | 46.26±0.93 |
| MCV (fL) | 47.10±0.45**** | 43.15±0.38 |
| MCH (pg) | 13.43±0.07**** | 12.56±0.10 |
| MCHC (g/dl) | 28.58±0.19 | 29.19±0.33 |
| RDW-SD (fl) | 41.85±1 | 42.76±2.21 |
| RET# (K/µl) | 735.05±37.73 | 646.62±141.29 |
| RET% (%) | 7.76±0.37 | 5.95±1.31 |
| PLT (K/µl) | 781.92±78.01 | 613.17±63.46 |
| WBC (K/µl) | 1.40±0.27 | 1.36±0.17 |
| NEUT# (K/µl) | 0.3±0.07 | 0.63±0.12 |
| LYMPH# (K/µl) | 0.93±0.18 | 0.58±0.09 |
| MONO# (K/µl) | 0.03±0 | 0.03±0.01 |
| EO# (K/µl) | 0.03±0.01 | 0.04±0.02 |
| BASO# | 0±0 | 0.01±0 |
| NEUT (%) | 23.58±1.44 | 48.16±4.91*** |
| LYMPH% (%) | 71.24±1.75*** | 46.47±4.39 |
| MONO % (%) | 2.77±0.3 | 2.77±0.33 |
| EO % (%) | 2.07±0.4 | 2.63±0.72 |
| BASO% (%) | 0.34±0.2 | 0.46±0.2 |
| RET-He (pg) | 17.86±0.19 | 17.02±0.23 |

Blood of CC037*ΔF508*/*ΔF508* and *wt/wt* was collected from the retro-orbital plexus at PND28 as described in material and methods. Hematologic parameters were evaluated using an automated cell counter (ProCyte Dx, IDEXX Laboratories, USA). Data are expressed as mean values ± standard errors of the means (SEM). Data represents the following genotypes and genders: *n*=13 *wt/wt* and *n*=12 *ΔF508/ΔF508* mice including number of males: *wt/wt* =7, *ΔF508/ΔF508*=7 and number of females: *wt/wt*=6*, ΔF508/ΔF508*=5. Statistical significance, determined by Mann-Whitney test non parametric test two-tailed, is indicated as follows: *, p<0.05; ****, p<0.0001.
