## Supplementary figures and images for "*ΔF508-Cftr* mutation in genetically diverse Collaborative Cross mice yields novel disease-relevant phenotypes for cystic fibrosis"

### Supplementary Figure 1

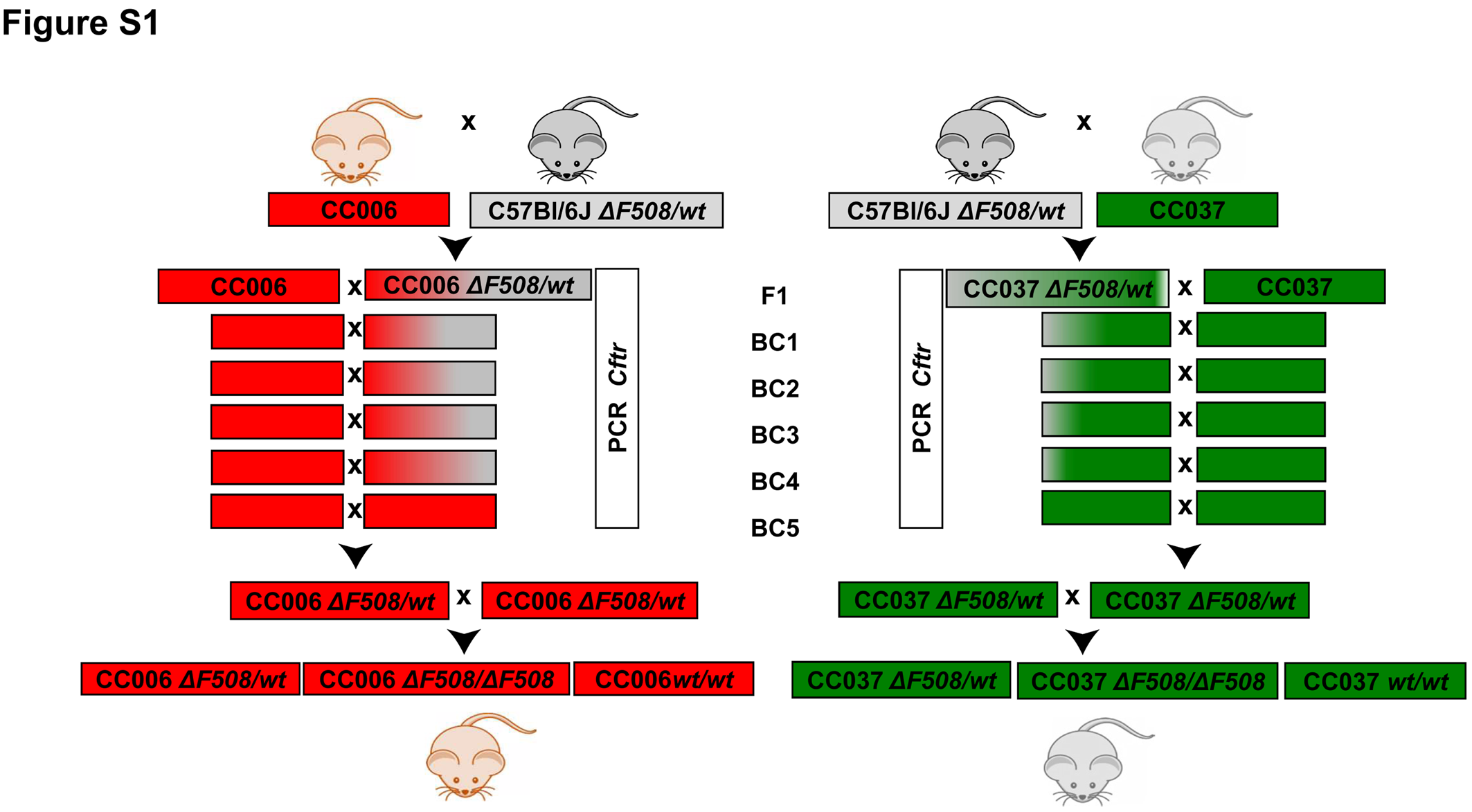

### Supplementary Figure 2

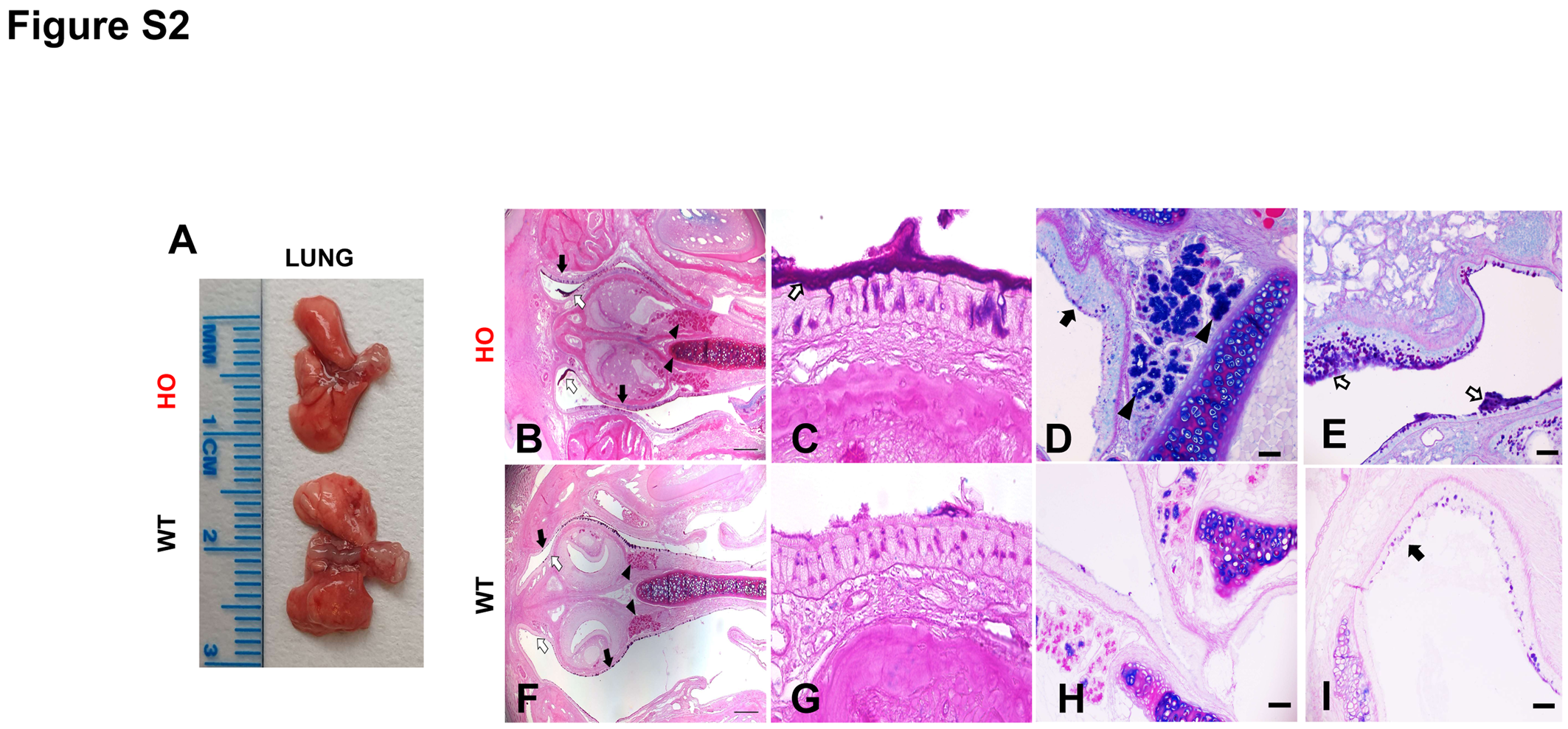

### Supplementary Figure 3

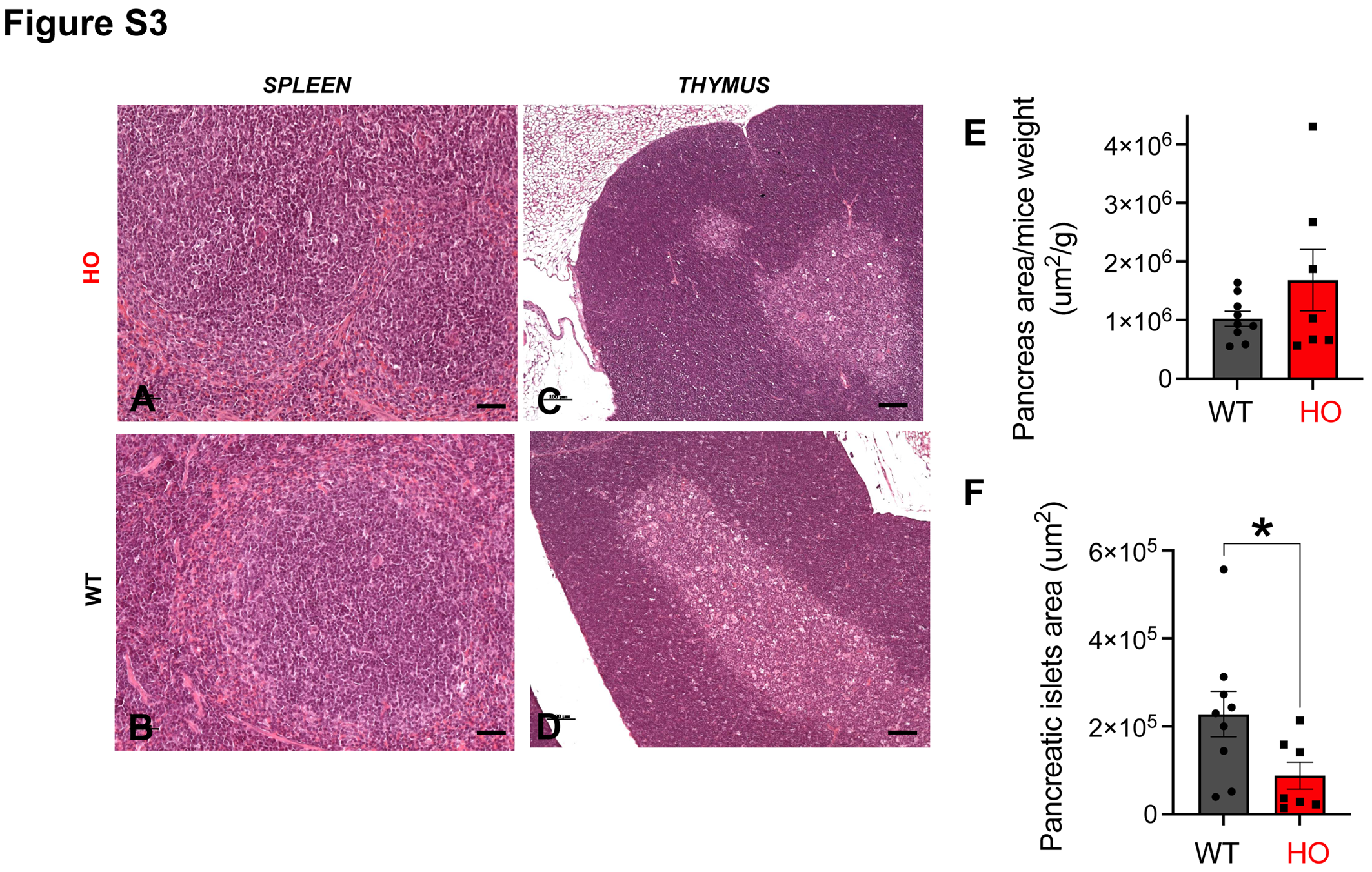
